# Increasing objective cardiometabolic burden associated with attenuations in the P3b event-related potential component in older adults

**DOI:** 10.1101/634873

**Authors:** Hannah A.D. Keage, Daniel Feuerriegel, Danielle Greaves, Emma Tregoweth, Scott Coussens, Ashleigh E. Smith

## Abstract

Cardiometabolic diseases and risk factors increase the risk of late-life cognitive impairment and dementia, and have also been associated with detrimental grey and white matter changes. However the functional brain changes associated with cardiometabolic health in late-life are unclear. We sought to characterise these functional changes by recording event-related potentials (ERPs) during a n-back working memory task (0, 1 and 2 back) in 85 adults (60% female) between 50 and 80 years of age. Due to a stratified recruitment approach, participants varied widely regarding cognitive function and cardiometabolic health. Standard and objective cut-offs for high blood glucose, waist to hip ratio (i.e. obesity), high blood cholesterol, and hypertension were employed to generate a summative score for cardiometabolic burden (none, one, or two or more above cut-off). Mixed effects modelling (covarying for age and gender) revealed no statistically significant associations between cardiometabolic burden and visual P1 and N1 component amplitudes. There was a significant effect for the P3b component: as cardiometabolic burden increased, P3b amplitude decreased. We show that cardiometabolic factors related to the development of cognitive impairment and dementia in late-life associate with functional brain activity, as recorded via ERPs. Findings have relevance for the monitoring of lifestyle interventions (typically targeting cardiometabolic factors) in ageing, as ERPs may provide a more sensitive measure of change than cognitive performance. Further, our results raise questions related to the findings of a broad range of ERP studies where the groups compared may differ in their cardiometabolic health status (not only in psychological symptomatology).

## Introduction

Cardiometabolic diseases and risk factors increase the risk of late-life cognitive impairment, including dementia (Livingston, et al., 2017; Norton, Matthews, Barnes, Yaffe, & Brayne, 2014). For example, Type 2 diabetes, obesity, physical inactivity, hypertension and high cholesterol all increase dementia risk. The greatest dementia risk is conveyed if these cardiometabolic factors are present in mid-life and early late-life, with null or paradoxical relationships seen in the oldest old, typically defined as those 85 years and over (Beydoun, Beydoun, & Wang, 2008; Harrison, et al., 2015). How these cardiometabolic factors affect brain structure and function between mid and late-life is the focus of current research, as it has been established that dementia-related pathologies accumulate decades before clinical symptoms (Jack Jr, et al., 2013). Such knowledge will enable us to understand the underlying neurophysiology of cardiometabolic-related late-life dementia risk.

Cardiometabolic factors, independently (Brundel, van den Heuvel, de Bresser, Kappelle, & Biessels; Chen, et al., 2015; Last, et al., 2007; Leritz, et al., 2011; Veit, et al., 2014; Walsh, Shaw, Sachdev, Anstey, & Cherbuin, 2017) and in combination (Fawns-Ritchie, et al., 2019; Tchistiakova & MacIntosh, 2016), have been reported to correlate with global and regional reductions in thickness and brain volume; along with more rapid cortical thinning over time. As compared to controls, older adults with Type 2 diabetes have lower global grey and white matter volumes (Walsh, Shaw, Sachdev, Anstey, & Cherbuin, 2017); reduced frontal white matter and parieto-occipital gray matter volumes (Last, et al., 2007); reduced right hemispheric cortical surface and volume (Brundel, van den Heuvel, de Bresser, Kappelle, & Biessels); and increased cortical thinning within the middle temporal gyrus, posterior cingulate gyrus, precuneus, entorhinal cortex, and right lateral occipital gyrus (Chen, et al., 2015). In young to mid-adulthood, widespread cortical thinning has been shown to relate independently to two obesity measures; body mass index (BMI) and visceral adipose tissue (Veit, et al., 2014).

Tchistiakova and MacIntosh (2016) combined cardiometabolic factors into a summative index (selection into study based on having one, two or three factors), from diabetes, smoking, blood pressure, fasting blood glucose and APOE genotype, in older adults from the Alzheimer’s Disease Neuroimaging Initiative (ADNI) with and without Mild Cognitive Impairment (MCI). Many factors (e.g. blood pressure) were extracted from medical records (including self-reported medication lists), rather than objectively; and notably, this study did not look at the effects of having no risk factors (i.e. participant selection was based on having at least one factor). Increases in the summative cardiometabolic factor score were associated with thinning of the temporal and frontal cortices (predominantly within the right hemisphere) in the MCI group, but not the control group (Tchistiakova & MacIntosh, 2016). Assessing a larger age range (44-79) in the UK Biobank, Fawns-Ritchie, et al. (2019) reported that cardiometabolic factors (smoking, hypertension, pulse pressure, diabetes, hypercholesterolaemia, body mass index, and waist–hip ratio) had detrimental and additive effects on the volumes of frontal and temporal cortex, subcortical structures, and white matter fibres (association and thalamic pathways). There appeared to be no hemispheric-bias in these results, and cognitive performance of the participants was not reported (Fawns-Ritchie, et al., 2019).

Poor cardiometabolic health has also been related to white matter structure, both cross-sectionally and longitudinally (Dufouil, et al., 2001; Fawns-Ritchie, et al., 2019; Fuhrmann, et al., 2019; Prins & Scheltens, 2015). A key and recent paper by Fuhrmann et al. (2019) demonstrated that blood pressure (lower diastolic and higher systolic), body mass (higher) and heart rate (higher) had independent detrimental effects on white matter macro- and micro-structure in a large cross-sectional population-based study from the UK.

In addition to structural brain changes, there is also evidence of functional brain changes associated with cardiometabolic risk factors for cognitive impairment and dementia (Braskie, Small, & Bookheimer, 2010; Chuang, et al., 2014). In a sample of older adults, Chuang, et al. (2014) reported that those with a higher cardiometabolic risk factor score (based on the Framingham risk assessment; Wolf, D’Agostino, Belanger, & Kannel, 1991) had greater task-related activation (fMRI) within the left inferior parietal region during an executive function task (adapted Flanker task); and that this activation related to poorer performance. Notably, no individual cardiometabolic factor was significantly associated with activation in this region (i.e. the effect was only apparent when factors were combined). Braskie, Small, and Bookheimer (2010) utilised a verbal paired associates learning task, and reported that in older cognitively healthy adults, higher cardiometabolic risk (summative score of BMI, systolic blood pressure and total cholesterol; with a focus on the former two) was associated with increased activation (as indexed using fMRI) within a large network including the posterior cingulate cortex, frontal, temporal and parietal regions; these associations held when controlling for the presence of the APOE ε4 allele (the major genetic risk allele for late-life dementia).

There appears to be little research using electroencephalography (EEG) to study the effects of cardiometabolic factors (associated with cognitive decline and dementia) on brain function. EEG has comparatively high temporal resolution compared to fMRI and enables insights into how cardiometabolic burden impacts different stages of sensory, perceptual and decision making processes. Event-related potentials (ERPs) are epochs of EEG data time-locked to a stimulus, typically averaged over multiple presentations of the same stimulus. ERPs enable us to index the time-course of critical processes for everyday cognition, including visual perception, categorisation and memory-related processes. We recorded ERPs while participants completed an executive function task, as performance in this domain appears particularly susceptible to cardiometabolic risk factors (Alcorn, et al., 2017; Chuang, et al., 2014; Elias, Elias, Robbins, & Budge, 2004), and has been reported to be the earliest domain affected by cardiometabolic factors, with impairment in other cognitive domains becoming apparent as cardiometabolic and vascular diseases progress (Fontbonne, Berr, Ducimetière, & Alpérovitch, 2001; Vasquez & Zakzanis, 2015). We also took the summative cardiometabolic factor approach (similar to Fawns-Ritchie, et al., 2019; Tchistiakova & MacIntosh, 2016), given that individual risk factors rarely occur in isolation.

Our primary aim was to determine whether and how objective cardiometabolic burden is associated with ERP component amplitudes during an executive function task; and whether associations were larger for broadly-distributed ERP components such as the P3b, as compared to more localised ERP components such as the visual P1 and N1. We employed the widely used n-back task to evoke the P1, N1 and P3b components, so that we could measure the effects of cardiometabolic burden on their amplitudes. This ERP-based approach enabled us to determine which components (each associated with different perceptual and cognitive processes) are attenuated, which is a critical and missing link between cardiometabolic burden and alterations in brain structure and performance. We hypothesised that smaller ERP component (P1, N1 and P3b) amplitudes would be associated with increasing cardiometabolic burden. A secondary aim investigated through exploratory analyses was to investigate if these cardiometabolic effects were dependent on hemisphere and cognitive impairment, as reported by Tchistiakova and MacIntosh (2016) using structural MRI.

## Methods

### Participants

A total of 88 adults (59% female) between 50 and 80 years of age completed the study. Participants were selectively recruited based on their self-reported cardiometabolic burden, so to gain a broad distribution of burden scores. Three participants were excluded from these analyses due to poor performance on the n-back tasks (greater than three standard deviations below the mean for either of both targets and non-targets; and/or below chance when assessing performance across targets and non-targets). Therefore the total number of participants in these analyses was 85, with 51 being female (60%). The mean age was 65.1 years (SD=7.5). Over half (54%) of participants were classified as having MCI or dementia according to the cut-offs of ≤92 on the Addenbrooke’s Cognitive Examination III (ACE-III) (Mioshi, Dawson, Mitchell, Arnold, & Hodges, 2006). ACE-III scores ranged between 72 and 99.

### Procedure

The study was approved by the University of South Australia Human Ethics Committee. Males and females aged 50 to 80 years were recruited if they self-reported as either low or high cardiovascular disease risk using the online Framingham risk assessment (Harrison, et al., 2017; Wolf, D’Agostino, Belanger, & Kannel, 1991). Within each decade age range of 50-59, 60-69, and 70-79 years we intended to recruit 30 participants (15 self-reported low cardiovascular disease risk and 15 self-reported high risk). Exclusion criteria were: history of stroke, clinical dementia diagnosis, blindness or vision problems not corrected by glasses/contact lenses, current diagnosis of a psychiatric disorder, an episode of unconsciousness for more than 5 minutes, and any known intellectual disabilities. Participants attended two three-hour appointments separated by approximately 8-10 days. During session one, informed consent, general health, cognition, fasted blood tests (minimum 8-hour fast), anthropometric assessments, blood pressure (arterial compliance measurement), and dietary assessments were conducted. EEG data was recorded during the second session.

### Cognitive performance

The ACE-III is a measure of cognitive functioning, and can be used to screen for cognitive impairment and dementia in older adults (Hsieh, Schubert, Hoon, Mioshi, & Hodges, 2013). The ACE-III consists of five subscales assessing attention/orientation (18 points), memory (26 points), fluency (14 points), language (26 points), and visuospatial abilities (16 points); therefore possible score range of 0 to 100. The test takes approximately 15-20 minutes to administer, and has demonstrated (along with its predecessor, the ACE-Revised) very good reliability, with an alpha coefficient of 0.8 (Hsieh, Schubert, Hoon, Mioshi, & Hodges, 2013; Mioshi, Dawson, Mitchell, Arnold, & Hodges, 2006). An overall ACE-III score is summed from the subtests, with higher scores indicating better cognitive function.

### Cardiometabolic burden

Blood glucose, waist to hip ratio (obesity), total blood cholesterol, and blood pressure were used as our cardiometabolic burden measures. Although there are various cut-offs reported across the previous literature, we used the most frequently reported. Participants were classified as positive for each cardiometabolic factor based on the following cut-offs: blood glucose ≥6.5 mmol/L (Carson, Reynolds, Fonseca, & Muntner, 2010; Howe-Davies, Simpson, & Turner, 1980); waist to hip ratio ≥.95 for men or ≥.90 for women (Gill, et al., 2003; WHO, 2011); total blood cholesterol ≥5.5 mmol/L (Solomon, et al., 2009); and if either diastolic blood pressure (BP) was ≥90 mmHg or systolic blood pressure was ≥140 mmHg (Muntner, et al., 2018; Whelton, et al., 2018).

Waist to hip ratios were obtained by research assistants trained in the standard protocols used by the International Society for the Advancement of Kinanthropometry. Using a luftkin executive thin line 2mm metal tape measure, waist measures were taken at the point of visible narrowing between the 10^th^ rib and the crest of the ilium during normal expiration. In the event there was no narrowing the measurement was taken at the mid-point between the lower costal (10^th^ rib) border and iliac crest. Hip measures were taken at the greatest point of posterior protuberance of the buttocks. Two separate measures were taken and if the measures differed by >20% a third measure was taken.

Participants rested horizontally for a minimum of five minutes in a dark room prior to blood pressure measurements. Blood pressure was non-invasively measured in conjunction with arterial compliance using a cardiovascular profiler (HDI cardiovascular profiler CR 2000, Hypertension Diagnostics, Minnesota, United States). The blood pressure cuff was fitted over the left brachial artery. Three readings were performed at 5 minute intervals and the average reading was calculated. If readings differed >20%, an additional reading was completed.

Approximately 23 mL of whole blood was collected via venepuncture into 2 x 9mL ethylenediaminetraacetic acid (EDTA) (18mg) anticoagulant and 1 x 4mL sodium fluoride Vacuette tubes (grenier bio-one, Kremsmünster, Austria). Following plasma separation at 4000 rpm for 10 minutes and frozen initially at −20°C for up to one week before transferred to −80°C until analysis samples were aliquoted into Eppendorf tubes and stored at −80 Celsuis until analysis. Cholesterol, triglycerides, HDL and glucose (from serum sample) were analysed in duplicate with a commercial assay kit (including quality controls and calibrators) using the KONELAB 20XTi (ThermoFisher, Massachusetts, United States).

### N-back tasks

Stimuli consisted of five capital letters (F, H, L, N, and T) presented in white font on a black background, subtending a visual angle of 1.4° (width) by 1.5° (height). For each n-back task, 50 of the stimuli were targets, and 100 were non-targets. Participants were seated comfortably 60cm in front of a computer monitor. Stimuli were presented in a pseudorandomised order one at a time for 500ms, followed by a blank screen. The inter-stimulus interval was jittered from 1200ms to 1500ms. Participants completed 0-, 1-, and 2-back versions of the task. Each n-back task lasted approximately 5 minutes.

For the 0-back task, participants were instructed to respond to the target letter (‘L’) with one hand (counterbalanced across participants). The 1-back task required participants to respond to a target whenever the letter presented matched the one presented immediately before it (i.e., the stimulus that was ‘one back’). For instance, if presented with the stimuli N-F-T-T, the participant would respond to the first three letters as non-targets, and the second ‘T’ as a target. Participants responded to target letters with one hand, and non-target letters with their other hand. The 2-back task required participants to respond to a target whenever the letter presented matched the one presented two trials previously. For example, if presented with the letters L-H-N-F-N, the participant would respond to the second ‘N’ as a target, and all other letters as non-targets. As with the other n-back tasks, target letters were responded to with one hand, and non-target letters with the opposite hand. For all n-back tasks, the first three stimuli were never targets and there were never more than two targets or more than two of the same stimulus presented in consecutive trials.

A short practice session was included before commencing each n-back task. Participants undertook one of two versions of each n-back task, which varied in trial order. The task version used was counterbalanced across participants. Response speed and accuracy were equally emphasised. Response hands assigned to targets and non-target response keys for responses was counterbalanced across participants.

### EEG recording and processing

We recorded EEG from 25 (Fp1, Fp2, AFpz, Fz, F3, F7, T7, C3, Cz, Pz, P3, P7, PO7, PO3, O1, Oz, O2, PO4, PO8, P8, P4, C4, T8, F8, F4) active electrodes using a Biosemi Active Two system (Biosemi, the Netherlands). Recordings were grounded using common mode sense and driven right leg electrodes (http://www.biosemi.com/faq/cms&drl.htm). EEG was sampled at 1024Hz (DC-coupled with an anti-aliasing filter, −3dB at 204Hz). Electrode offsets were kept within ±50μV. We processed EEG data using EEGLab V.13.4.4b (Delorme & Makeig, 2004) and ERPLab V.4.0.3.1 (Lopez-Calderon and Luck, 2014) running in MATLAB (The Mathworks).

Data were resampled to 512Hz and re-referenced offline to a nose reference. Bad sections of EEG data (e.g., containing large or atypical artefacts) were removed manually. Excessively noisy channels were identified by visual inspection and were not included as input data for the independent components analysis (ICA). 50 Hz line noise was identified using Cleanline (Mullen, 2012) using a separate 1 Hz high-pass filtered dataset (EEGLab Basic FIR Filter New, zero-phase, finite impulse response, −6 dB cutoff frequency 0.5 Hz, transition bandwidth 1 Hz). Identified line noise was subtracted from the unfiltered dataset (as recommended by Bigdely-Shamlo, Mullen, Kothe, Su, & Robbins, 2015). A separate dataset was processed in the same way, except a 1 Hz high-pass filter was applied (filter settings as above) to improve stationarity for the ICA (as done by Feuerriegel, Churches, Coussens, & Keage, 2018). ICA was performed on the 1 Hz high-pass filtered dataset (RunICA extended algorithm, Jung, et al., 2000). Independent component information was transferred to the unfiltered dataset. Independent components associated with ocular activity (i.e., blinks and saccades) were identified and removed according to guidelines in Chaumon, Bishop, and Busch (2015).

Bad channels were interpolated using the cleaned data. The dataset were then high-pass filtered at 0.1Hz and low-pass filtered at 30Hz (EEGLab FIR Filter New, default transition band widths). Data was epoched from −100 to +800ms. Epochs containing amplitudes larger than ±100 μV in any channel were excluded from analyses. ERPs were then averaged across epochs according to target/non-target status and participant response (hit/miss/correct rejection/false alarm) for correct trials only (i.e. error trials excluded).

We then calculated the mean amplitudes for the P1, N1 and P3b ERP components using the following time windows: 80-120ms for the visual P1, 120-180ms for the visual N1, and 300-550ms for the P3b. Only trials with correct rejections of non-target stimuli were included, as we had no hypotheses related to target detection processes, and this condition had more trials for averaging (therefore reducing noise). Further, the P3a component is also prominent following target stimuli, which is largest over frontocentral sites (rather than posterior channels for the P3b), which was another reason for the exclusion of target trial ERPs from analyses. For the P1 and N1 components the following electrodes were included (averaged relative to hemisphere): P7, PO7 and O1 (left hemisphere), and P8, PO8 and O2 (right hemisphere). For the P3b components the P3 and C3 electrodes were averaged for the left hemisphere, and P4 and C4 for the right hemisphere.

### Statistical Approach

STATA v15.1 IC was used for all analyses. Behavioural data during the n-back tasks were analysed using three separate mixed effects models with maximum likelihood estimation and ID set as both a random intercept and slope, with the following outcomes: (1) reaction time to correct hits to targets, (2) accuracy of hits to targets, and (3), accuracy of correct non-responses to non-targets. For each of these models, difficulty (0-, 1-, and 2-back), cardiometabolic burden (0=none, 1=one, 2=two or factors above cut-off), along with an interaction between difficulty and cardiometabolic burden were used as predictors.

Mixed effects modelling with maximum likelihood estimation, with ID set as both a random intercept and slope, was conducted for each component mean amplitude as outcomes: P1, N1 and P3. Age and gender were used as covariates. Cardiometabolic burden (0=none, 1=one, 2=two or factors above cut-off) was used as the predictor variable, which enabled us to address our hypotheses; with the other predictors being hemisphere (left and right hemisphere electrodes) and difficulty (0, 1 and 2-back tasks). We included hemisphere and difficulty to make parallels with previous results, given the n-back task has been widely used, and further, to compare effect sizes with cardiometabolic burden. We ran two sets of follow-up exploratory stratified analyses, which were stratified by (1) hemisphere (left and right as separate models) and (2) cognitive impairment (those ≤92 on the ACE-III); in line with structural MRI findings from Tchistiakova and MacIntosh (2016). To account for multiple comparisons we lowered our critical alpha value to .017 (.05 divided by 3), taking into account that we assessed three ERP components (P1, N1 and P3). Cohen *f^2^* was our measure of effect size, with *f^2^* values > .02, >.15, and >.35 representing small, medium, and large effect sizes, respectively.

## Results

### Distribution of cardiometabolic burden

The distribution of cardiometabolic burden is displayed in Table 1, relative to each of the four cardiometabolic factors cut-offs, and the summative variable (number of factors above the cut-off: 0, 1 or 2 or more). Our participant sampling strategy resulted in a roughly equivalent number of participants with 0, 1 and 2 or more risk factors.

**Table 1.** Distribution of cardiometabolic burden across sample.

| | High blood glucose<br>($\geq 6.5$ mmol/L) | High waist:hip<br>($\geq .95$ men/<br>$\geq .90$ women) | High total blood<br>cholesterol<br>($\geq 5.5$ mmol/L) | Hypertension<br>(diastolic $\geq 90$/<br>systolic $\geq 140$<br>mmHg) | Summative<br>cardiometabolic burden | | |
| --- | --- | --- | --- | --- | --- | --- | --- |
| | | | | | 0 | 1 | $\geq 2$ |
| Women<br>(n=51) | 9<br>(18%) | 9<br>(18%) | 20<br>(39%) | 14<br>(27%) | 15<br>(29%) | 23<br>(45%) | 13<br>(25%) |
| Men<br>(n=34) | 13<br>(38%) | 20<br>(59%) | 8<br>(24%) | 13<br>(38%) | 5<br>(15%) | 11<br>(32%) | 18<br>(53%) |
| Total<br>(n=85) | 22<br>(26%) | 29<br>(34%) | 28<br>(33%) | 27<br>(32%) | 20<br>(24%) | 34<br>(40%) | 31<br>(36%) |

### Performance on the n-back tasks relative to cardiometabolic burden

Performance on the n-back tasks across the cardiometabolic burden groups is displayed in Table 2. Mixed effects modelling revealed no significant effect of cardiometabolic burden (*p*=.922) nor interaction between cardiometabolic burden and difficulty (*p*=.719), on reaction times (to correct hits to targets). When the accuracy of target detection was used as the outcome, there was a significant interaction between vascular burden and difficulty (beta=−1.455, SE=0.700, z=−2.08, *p*=.037, 95%CI −2.827 to −0.085), which reflected that the number of correct hits to targets were lower in those with high cardiometaolic burden, but only when difficulty was high; there was no main effect of cardiometabolic burden (*p*=.207). This pattern of effects was mirrored when assessing the accuracy of correct rejections of non-targets, however the effect for the interaction missed conventional significance (*p*=.096). Difficulty was a significant predictor in all three models, with RT and accuracy decreasing as difficulty increased (all *p*<.001).

**Table 2.** Performance (reaction time and accuracy) on the n-back tasks relative to cardiometabolic burden.

| | | No cardiometabolic burden | 1 cardiometabolic factor | $\geq 2$ cardiometabolic factor | Total |
| --- | --- | --- | --- | --- | --- |
| Reaction time |  |  |  |  |  |
| 0-back | mean | 411.39 | 423.59 | 425.25 | 421.54 |
|  | SD | 56.34 | 48.66 | 49.98 | 50.56 |
| 1-back | mean | 506.58 | 522.05 | 517.86 | 517.07 |
|  | SD | 98.10 | 92.51 | 65.24 | 83.73 |
| 2-back | mean | 710.16 | 737.31 | 737.40 | 731.38 |
|  | SD | 118.00 | 136.90 | 127.44 | 128.24 |
| Accuracy: correct hits (to targets) |  |  |  |  |  |
| 0-back | n trials | 49 | 49 | 49 | 49 |
|  | % | 98 | 98 | 98 | 98 |
| 1-back | n trials | 48 | 48 | 48 | 48 |
|  | % | 96 | 96 | 96 | 96 |
| 2-back | n trials | 32 | 27 | 26 | 28 |
|  | % | 64 | 54 | 52 | 56 |
| Accuracy: correct rejections (to non-targets) |  |  |  |  |  |
| 0-back | n trials | 97 | 97 | 97 | 97 |
|  | % | 97 | 97 | 97 | 97 |
| 1-back | n trials | 97 | 96 | 97 | 96 |
|  | % | 97 | 96 | 97 | 96 |
| 2-back | n trials | 89 | 86 | 85 | 86 |
|  | % | 89 | 86 | 85 | 86 |

### Associations between *cardiometabolic* burden and ERP component amplitude

Table 3 details all results from the primary mixed effect models. For the P1 component, the mixed model revealed a small effect for cardiometabolic burden (with smaller amplitudes as cardiometabolic burden increased), however this did not meet our adjusted alpha value (*p*=.037; alpha=.017). For the mixed models with N1 as the outcome there was an effect of difficulty (with smaller amplitudes for more difficult versions of the task). In this model no other effects were statistically significant. For the P3b component, the mixed effects model revealed significant main effects for both hemisphere and cardiometabolic burden; with the P3b component larger over the right hemisphere, and smaller P3b components evoked in participants with higher cardiometabolic burden. For each one unit increase in cardiometabolic burden (i.e. 0 to 1, and 1 to ≥2) amplitude of the P3b decreased by 0.703μV (in the context of the P3b having a mean of 1.320 μV and SD of 2.078 across all conditions). Notably, the effect size for cardiometabolic burden was small (*f^2^* <.001). Grand average ERPs for each cardiometabolic burden group are displayed in Figure 1.

**Table 3.** Results from mixed effects models for the P1, N1 and P3b ERP component amplitudes (primary analyses).

|  | Beta | Standard Error | z | p | 95%CI |  | f<sup>2</sup> |
| --- | --- | --- | --- | --- | --- | --- | --- |
| *P1 component* |  |  |  |  |  |  |  |
| Age | 0.021 | 0.027 | 0.80 | .422 | -0.031 | 0.073 | <.001 |
| Gender* | -0.353 | 0.419 | -0.84 | .399 | -1.175 | 0.468 | <.001 |
| Difficulty | 0.052 | 0.060 | 0.88 | .379 | -0.065 | 0.171 | .002 |
| Hemisphere* | 0.171 | 0.098 | 1.74 | .081 | 0.021 | 0.364 | .008 |
| Cardiometabolic burden | -0.562 | 0.270 | -2.08 | .037 | -1.090 | -0.033 | <.001 |
| *N1 component* |  |  |  |  |  |  |  |
| Age | -0.005 | 0.039 | -0.14 | .890 | -0.082 | 0.071 | <.001 |
| Gender* | 0.431 | 0.614 | 0.70 | .482 | -0.772 | 1.635 | <.001 |
| Difficulty | 0.534 | 0.064 | 8.33 | <.001 | 0.409 | 0.660 | .170 |
| Hemisphere* | -0.101 | 0.105 | -0.96 | .336 | -0.307 | 0.105 | .002 |
| Cardiometabolic burden | 0.136 | 0.395 | 0.34 | .731 | -0.638 | 0.909 | <.001 |
| *P3b component* |  |  |  |  |  |  |  |
| Age | 0.029 | 0.026 | 1.13 | .260 | -0.022 | 0.080 | <.001 |
| Gender* | -0.170 | 0.412 | -0.41 | .680 | -0.976 | 0.637 | <.001 |
| Difficulty | 0.056 | 0.050 | 1.11 | .266 | -0.043 | 0.155 | .001 |
| Hemisphere* | 0.294 | 0.082 | 3.58 | <.001 | 0.133 | 0.455 | .014 |
| Cardiometabolic burden | -0.703 | 0.265 | -2.65 | .008 | -1.223 | -0.184 | <.001 |

\*Gender = 1 female, 2 male; Hemisphere = 1 left, 2 right.

**Figure 1.**
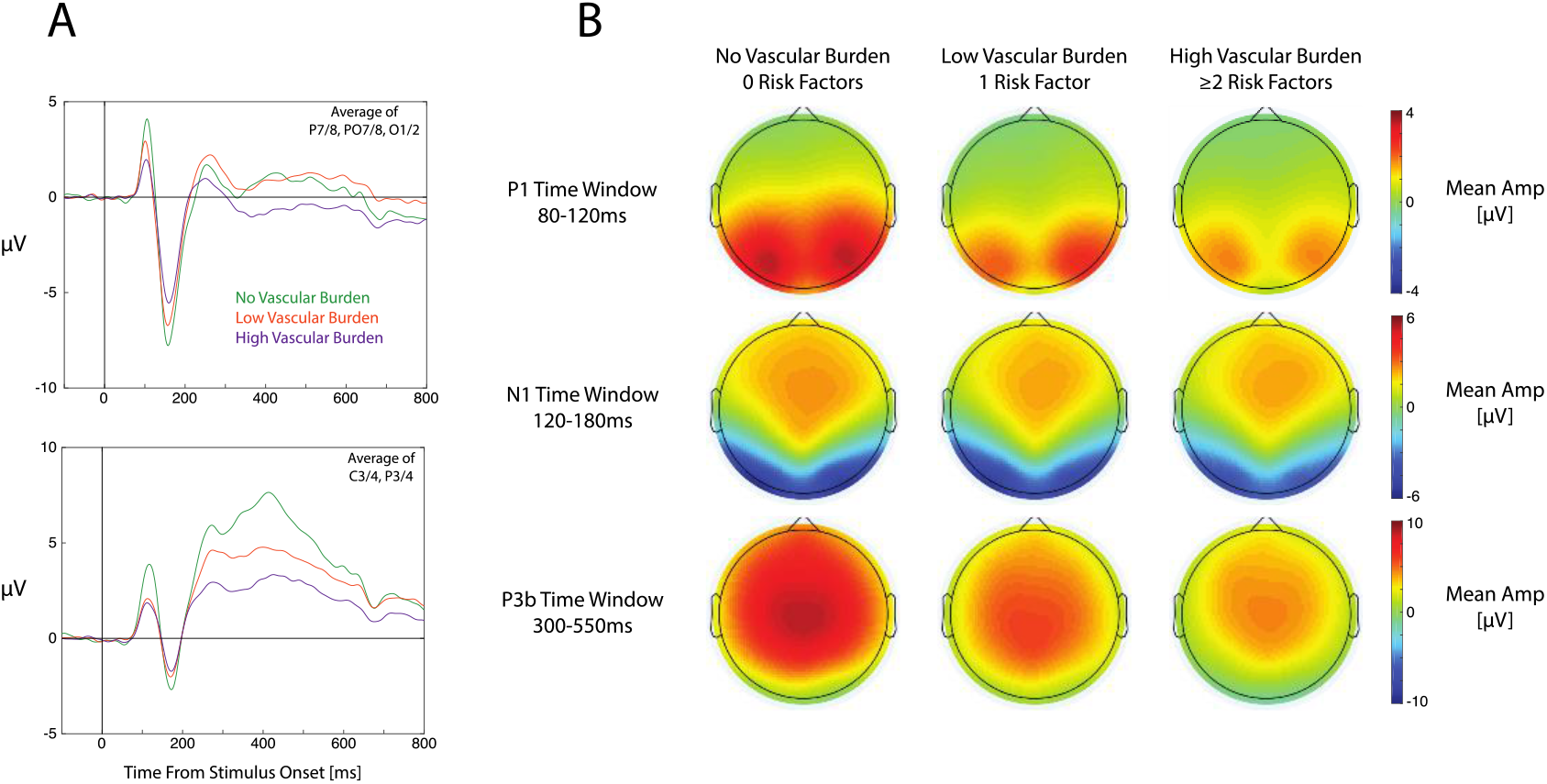
Grand average ERPs and topographic plots for each cardiometabolic burden group. A) Grand average ERP waveforms for each group averaged over electrodes P7/8, PO7/8 and O1/2 (top panel) and averaged over C3/4 and P3/4 (bottom panel). ERPs were also averaged across versions of the n-back task in these plots. B) Topographic maps displaying mean amplitudes for each group during the measurement windows for the visual P1 (top row), visual N1 (middle row) and P3b (bottom row).

Secondary analyses assessed effects stratified by hemisphere (results displayed in Table 3) and cognitive impairment status (results displayed in Table 4). The observed pattern of effects did not appear to differ between the hemispheres. Effects did appear to differ between cognitive impairment groups, with cardiometabolic burden displaying significant negative effects for both P1 and P3b amplitudes within the cognitively impaired group only (both with a small effect size). Notably, we took this stratified approach in line with Tchistiakova and MacIntosh (2016). An alternative would be including interaction terms (with hemisphere and cognitive impairment status respectively); when we did do this, interaction effects were no longer statistically significant, due to the small effect sizes visible in the stratified analyses.

**Table 4.** Results from mixed effects models for the P1, N1 and P3b ERP component amplitudes stratified by hemisphere (secondary analyses).

|  | Beta | Standard Error | *z* | *p* | 95%CI |  | *f*<sup>2</sup> |
| --- | --- | --- | --- | --- | --- | --- | --- |
| *P1 component: left hemisphere* |  |  |  |  |  |  |  |
| Age | 0.015 | 0.028 | 0.54 | .586 | -0.039 | 0.069 | <.001 |
| Gender* | -0.251 | 0.437 | -0.57 | .566 | -1.106 | 0.605 | <.001 |
| Difficulty | 0.139 | 0.094 | 1.49 | .137 | -0.044 | 0.323 | .014 |
| Cardiometabolic burden | -0.484 | 0.281 | -1.72 | .085 | -1.035 | -0.066 | <.001 |
| *P1 component: right hemisphere* |  |  |  |  |  |  |  |
| Age | 0.028 | 0.026 | 1.09 | .276 | -0.023 | 0.080 | <.001 |
| Gender* | -0.493 | 0.413 | -1.19 | .233 | -1.304 | 0.317 | <.001 |
| Difficulty | -0.025 | 0.080 | -0.31 | .759 | -1.181 | 0.132 | .001 |
| Cardiometabolic burden | -0.631 | 0.266 | -2.38 | .017 | -1.152 | -0.111 | <.001 |
| *N1 component: left hemisphere* |  |  |  |  |  |  |  |
| Age | -0.014 | 0.038 | -0.37 | .710 | -0.090 | 0.062 | <.001 |
| Gender* | 0.469 | 0.612 | 0.77 | .444 | -0.731 | 1.667 | <.001 |
| Difficulty | 0.484 | 0.105 | 4.63 | <.001 | 0.279 | 0.690 | .131 |
| Cardiometabolic burden | 0.206 | 0.394 | 0.52 | .600 | -0.565 | 0.978 | <.001 |
| *N1 component: right hemisphere* |  |  |  |  |  |  |  |
| Age | 0.002 | 0.040 | 0.06 | .952 | -0.076 | 0.080 | <.001 |
| Gender* | 0.388 | 0.628 | 0.62 | .536 | -0.842 | 1.618 | <.001 |
| Difficulty | 0.584 | 0.084 | 6.96 | <.001 | 0.419 | 0.748 | .297 |
| Cardiometabolic burden | 0.057 | 0.402 | 0.14 | .887 | -0.730 | 0.845 | <.001 |
| *P3b component: left hemisphere* |  |  |  |  |  |  |  |
| Age | 0.026 | 0.027 | 0.96 | .335 | -0.027 | 0.078 | <.001 |
| Gender* | -0.037 | 0.421 | -0.09 | .930 | -0.861 | 0.787 | <.001 |
| Difficulty | 0.195 | 0.074 | 2.64 | .008 | 0.050 | 0.340 | .017 |
| Cardiometabolic burden | -0.670 | 0.271 | -2.48 | .013 | -1.201 | -0.140 | <.001 |
| *P3b component: right hemisphere* |  |  |  |  |  |  |  |
| Age | 0.033 | 0.027 | 1.23 | .220 | -0.020 | 0.086 | <.001 |
| Gender* | -0.302 | 0.425 | -0.71 | .477 | -1.135 | 0.530 | <.001 |
| Difficulty | -0.083 | 0.066 | -1.26 | .208 | -0.212 | 0.046 | .004 |
| Cardiometabolic burden | -0.736 | 0.274 | -2.67 | .007 | -1.272 | -0.200 | <.001 |

\*Gender = 1 female, 2 male

**Table 5.** Results from mixed effects models for the P1, N1 and P3b ERP component amplitudes stratified by cognitive status (secondary analyses).

|  | Beta | Standard Error | z | p | 95%CI |  | f<sup>2</sup> |
| --- | --- | --- | --- | --- | --- | --- | --- |
| *P1 component: no cognitive impairment* |  |  |  |  |  |  |  |
| Age | 0.011 | 0.047 | 0.22 | .823 | -0.082 | 0.103 | <.001 |
| Gender* | 0.093 | 0.663 | 0.14 | .888 | -1.207 | 1.394 | <.001 |
| Difficulty | -0.139 | 0.092 | -1.51 | .131 | -0.320 | 0.041 | .013 |
| Hemisphere* | 0.251 | 0.151 | 1.66 | .097 | -0.045 | 0.548 | .016 |
| Cardiometabolic burden | -0.119 | 0.407 | -0.29 | .771 | -0.916 | 0.679 | <.001 |
| *P1 component: cognitive impairment* |  |  |  |  |  |  |  |
| Age | 0.027 | 0.031 | 0.87 | .382 | -0.033 | 0.087 | <.001 |
| Gender* | -0.606 | 0.516 | -1.17 | .240 | -1.617 | 0.405 | <.001 |
| Difficulty | 0.208 | 0.077 | 2.69 | .007 | 0.057 | 0.360 | .032 |
| Hemisphere* | 0.111 | 0.126 | 0.88 | .380 | -0.137 | 0.358 | .003 |
| Cardiometabolic burden | -0.901 | 0.345 | -2.61 | .009 | -1.578 | -0.224 | <.001 |
| *N1 component: no cognitive impairment* |  |  |  |  |  |  |  |
| Age | 0.009 | 0.065 | 0.14 | .892 | -0.118 | 0.134 | <.001 |
| Gender* | -0.007 | 0.910 | -0.01 | .994 | -1.791 | 1.777 | <.001 |
| Difficulty | 0.593 | 0.093 | 6.35 | <.001 | 0.410 | 0.776 | .220 |
| Hemisphere* | -0.133 | 0.153 | -0.86 | .387 | -0.433 | 0.168 | .004 |
| Cardiometabolic burden | 0.363 | 0.563 | 0.65 | .518 | -0.739 | 1.466 | <.001 |
| *N1 component: cognitive impairment* |  |  |  |  |  |  |  |
| Age | -0.010 | 0.049 | -0.21 | .836 | -0.106 | 0.086 | <.001 |
| Gender* | 0.834 | 0.830 | 1.01 | .315 | -0.792 | 2.460 | <.001 |
| Difficulty | 0.486 | 0.088 | 5.53 | <.001 | 0.313 | 0.658 | .136 |
| Hemisphere* | -0.075 | 0.144 | -0.53 | .599 | -0.357 | 0.206 | .001 |
| Cardiometabolic burden | -0.118 | 0.553 | -0.21 | .831 | -1.202 | 0.966 | <.001 |
| *P3b component: no cognitive impairment* |  |  |  |  |  |  |  |
| Age | -0.020 | 0.050 | -0.40 | .691 | -0.118 | 0.078 | <.001 |
| Gender* | 0.560 | 0.701 | 0.80 | .424 | -0.813 | 1.933 | <.001 |
| Hemisphere* | -0.211 | 0.076 | -2.77 | .006 | -0.361 | -0.062 | .019 |
| Difficulty | 0.395 | 0.125 | 3.16 | .002 | 0.150 | 0.640 | .025 |
| Cardiometabolic burden | -0.365 | 0.433 | -0.84 | .399 | -1.214 | 0.484 | <.001 |
| *P3b component: cognitive impairment* |  |  |  |  |  |  |  |
| Age | 0.064 | 0.026 | 2.45 | .014 | 0.013 | 0.115 | <.001 |
| Gender* | -0.619 | 0.439 | -1.41 | .158 | -1.479 | 0.241 | <.001 |
| Difficulty | 0.276 | 0.065 | 4.24 | <.001 | 0.148 | 0.403 | .036 |
| Hemisphere* | 0.212 | 0.106 | 1.99 | .046 | 0.003 | 0.420 | .008 |
| Cardiometabolic burden | -0.880 | 0.294 | -3.00 | .003 | -1.455 | -0.304 | <.001 |

\*Gender = 1 female, 2 male; Hemisphere = 1 left, 2 right.

## Discussion

We show that objective cardiometabolic burden is associated with attenuations in P3b ERP component amplitudes during an executive function task. Although effects were in the same direction for the earlier P1 and N1 components (i.e. smaller components with increasing cardiometabolic burden), effects were small and not statistically significant. When those with and without a cognitive impairment were assessed separately, a similar effect on P3b amplitudes was apparent, along with an effect for the P1 component, in those with cognitive impairment. This pattern of effects is similar to a structural MRI study by Tchistiakova and MacIntosh (2016) that found cardiometabolic burden was associated with volume reductions only in those with MCI. However we did not find the right hemispheric predominance as reported by Tchistiakova and MacIntosh (2016).

The P3b is a large and broad component, seen across much of the scalp, whereas the earlier components are relatively small and localised over occipito-parietal regions, as is typical for visually presented stimuli. We propose two reasons for seeing a significant effect for the P3b, but not earlier components: (1) small effect sizes for cardiometabolic burden and (2), differences in the extent of neural generators. Small effect sizes for cardiometabolic burden on brain structure have been previously reported (Fawns-Ritchie, et al., 2019). Given the effect size of cardiometabolic burden was small (as seen here across all components), such effects would be preferentially found for larger amplitude ERP components that can be recorded with a high signal-to-noise ratio, such as the P3b. In terms of neural generators, the P1 and N1 predominately relate to activity in visual cortex and ventral temporal areas, whereas the candidate regions contributing to the P3b include frontal, parietal and tempo-occipital regions (e.g. Bachiller, et al., 2015; Volpe, et al., 2007) although the exact regions remain a topic of debate. Fawns-Ritchie, et al. (2019) reported that frontal and temporal cortical thinning is associated with cardiometabolic burden in a large UK sample; other studies have reported more widespread cortical thinning (Brundel, van den Heuvel, de Bresser, Kappelle, & Biessels; Chen, et al., 2015; Last, et al., 2007; Leritz, et al., 2011; Veit, et al., 2014; Walsh, Shaw, Sachdev, Anstey, & Cherbuin, 2017). Cumulative effects on multiple neural generators of the P3b is likely why widespread cortical thinning has larger effects on the P3b and smaller effects on more localised components such as the P1 and N1 components (at least when the whole group was included in analyses).

To our knowledge, no ERP study has assessed associations with a composite (or even multiple independent markers of) cardiometabolic health. Attenuated P3b component amplitudes have been reported in obese children (Tascilar, et al., 2011). However, many psychiatric conditions are associated with cardiometabolic burden, such as Schizophrenia, even around the time of diagnosis (Correll, et al., 2014). There are many decades of research assessing and describing ERP differences between clinical and control groups (Kappenman & Luck, 2016), including Schizophrenia (Feuerriegel, Churches, Hofmann, & Keage, 2015; McCleery, et al., 2015). From our data, it is likely that smaller ERP component amplitudes are not only due to experimental and paradigm factors such as impaired sensory coding, perceptual categorisation, and impaired attention; but rather, also physiological differences, related to cardiometabolic health and its related downstream cerebrovascular (Last, et al., 2007), structural grey and white changes (Fawns-Ritchie, et al., 2019), along with neurochemical and neuropathological associations (Kalaria, 2010). Differences between clinical and control groups in existing ERP studies may not be due entirely to cognitive deficits (for example in attention) but rather, at least in part, due to differences in cardiometabolic health.

Given the critical role ascribed the P3b in relation to accumulating sensory evidence in perceptual decisions (e.g. Twomey, Murphy, Kelly, & O’Connell, 2015), our results also suggest caution when comparing P3b amplitudes across groups with different cardiometabolic burden. For example, P3b peak amplitudes gradually decline from adolescence to older age (Rossini, Rossi, Babiloni, & Polich, 2007). Reductions in participants’ P3b amplitudes have been interpreted as requiring less evidence to reach a decision, yet computational modelling studies indicate that older adults are actually more conservative in their decision-making, requiring more evidence for a decision than younger adults (reviewed in Dully, McGovern, & O’Connell, 2018). This discrepancy between ERPs and patterns of behavioural results may be partly explained by structural and functional changes associated with cardiometabolic burden, as reported in our study.

The major limitation of our study is that it is cross-sectional. Recent structural brain imaging studies have demonstrated patterns of change associated with cardiometabolic health that are different (to some extent) to cross-sectional studies. Walsh, Shaw, Sachdev, Anstey, and Cherbuin (2017) recently demonstrated that elevated blood glucose was associated with reductions of global grey matter volume over four years. Over eight years of follow-up in older adults from the Baltimore Longitudinal Study of Aging, Gonzalez, Pacheco, Beason-Held, and Resnick (2015) showed an increased rate of thinning in several brain regions in hypertensive individuals, as compared with normotensive individuals, including the left frontomarginal gyrus in the left hemisphere and the right superior temporal, fusiform, and lateral orbitofrontal cortex. Higher midlife blood pressure and longer durations of hypertension were associated with accelerated rates cortical thinning in the right superior temporal gyrus (Gonzalez, Pacheco, Beason-Held, & Resnick, 2015). Future ERP studies should assess longitudinal effects of cardiometabolic health. Another limitation is that we could not include smoking, which is a major vascular-related risk factor for dementia (Livingston, et al., 2017), as we had only a small number of smokers in our sample (which meant that we could not reliably assess such effects).

The major strength of this study is the objective characterisation of cardiometabolic health, as self-reported health is not often accurate (Johnston, Propper, & Shields, 2009; Okura, Urban, Mahoney, Jacobsen, & Rodeheffer, 2004). In addition, our stratified sampling approach enabled us to have a large spread of cardiometabolic health. Lastly, we employed a widely-used cognitive paradigm (the n-back task), which permits the comparison of the strength of effects of cardiometabolic health against standard factors such as hemisphere and difficulty.

We have characterised cardiometabolic health objectively in a large sample of older adults, to demonstrate that these factors are associated with functional brain activity. Effects were most notable for the P3b component, where increasing number of cardiometabolic factors above standard cut-offs was associated with attenuations in the component; and most notable in those with cognitive impairment. Although effect sizes were small, they are of great importance. This study is the first to identify that functional brain health is influenced by broader cardiometabolic health, which of note, are modifiable risk factors for late-life dementia (Livingston, et al., 2017; Norton, Matthews, Barnes, Yaffe, & Brayne, 2014). ERPs may be more sensitive than cognitive tests to index intervention-related brain changes in late-life, for example, in multi-component or vascular interventions (Kivipelto, et al., 2013; Ngandu, et al., 2015; SPRINT-MIND-Investigators, 2019). Findings also have implications for the larger ERP literature, whereby cardiometabolic health should be considered as a factor influencing group differences (not just cognitive differences, such as attention).

